# *Seek-a-Seq*: Optical detection of target nucleic acids with attomolar sensitivity

**DOI:** 10.1101/2025.02.18.638747

**Authors:** Anand Kant Das, Alam Saj, Hin Hark Gan, Kristin C. Gunsalus, George T. Shubeita

## Abstract

We introduce Seek-a-Seq, an enzyme-free, PCR amplification-free, optical method to detect microbial nucleic acids based on fluorescent readout of hybridization between target sequences and complementary probes immobilized on the surface of customized sample chambers. We demonstrate the ability to detect ultralow concentration (~ 1 attomolar) of short target DNA and synthetic SARS-CoV-2 RNA with ease and specificity. Seek-a-Seq can enormously reduce costs, accelerate diagnostic workflows, and outperform existing enzyme-dependent methods.

---

Nucleic acid detection is central to disease diagnostics, genetic testing, infectious disease surveillance, and environmental monitoring^1,2^ Most existing methods rely on nucleic acid amplification tests (NAATs), such as polymerase chain reaction (PCR) and loop-mediated isothermal amplification (LAMP), to enhance detection sensitivity by amplifying microbial nucleic acids. However, NAATs require multistep workflows, complex reaction conditions, specialized reagents, dedicated equipment, and trained personnel, making them less suitable for rapid point-of-care deployment^3^. Additionally, preamplification increases the risk of contamination, false positives, reagent instability, and variability. In the event of a widespread disease outbreak or pandemic, the demand for specific enzymes and reagents surges, exacerbating supply chain challenges and limiting accessibility. Nonetheless, nucleic-acid based methods continue to evolve with advancements in technology, including the development of novel probes, detection platforms, and multiplexing strategies^4–11^. There is an urgent need for simple, sensitive, amplification-free nucleic acid detection technologies with multiplexing capabilities that can facilitate point-of-care applications, streamline diagnostic workflows, and significantly reduce costs^12^.

Here, we present an amplification-free, single-molecule optical technique for detecting specific nucleic acid sequences with extremely high sensitivity and specificity. Seek-a-Seq relies on a simple and versatile imaging platform capable of directly detecting a few hundred molecules of target nucleic acids in a milliliter of solution. The core principle is based on direct hybridization between complementary probe sequences immobilized on a functionalized glass surface and the target nucleic acid (**Figure 1A**). Subsequent exposure to cyanine-based intercalator dyes (e.g., SYBR Safe), followed by total internal reflection fluorescence (TIRF) imaging, generates a high signal-to-noise fluorescence output, confirming the presence of the target nucleic acid. The immobilization of short DNA probes on glass surfaces can introduce surface artifacts, nonspecific binding of fluorescent species, or unwanted DNA contamination, potentially leading to false-positive detections. To eliminate this, we optimized a surface and assay chamber passivation protocol (Methods, **Figure S1**). Biotinylated single-stranded DNA (ssDNA) probes were immobilized on the imaging surface via a biotin-PEG–streptavidin bridge (**Figure 1A**), leveraging the exceptionally strong, non-covalent interaction between streptavidin and biotin.

**Figure 1:**
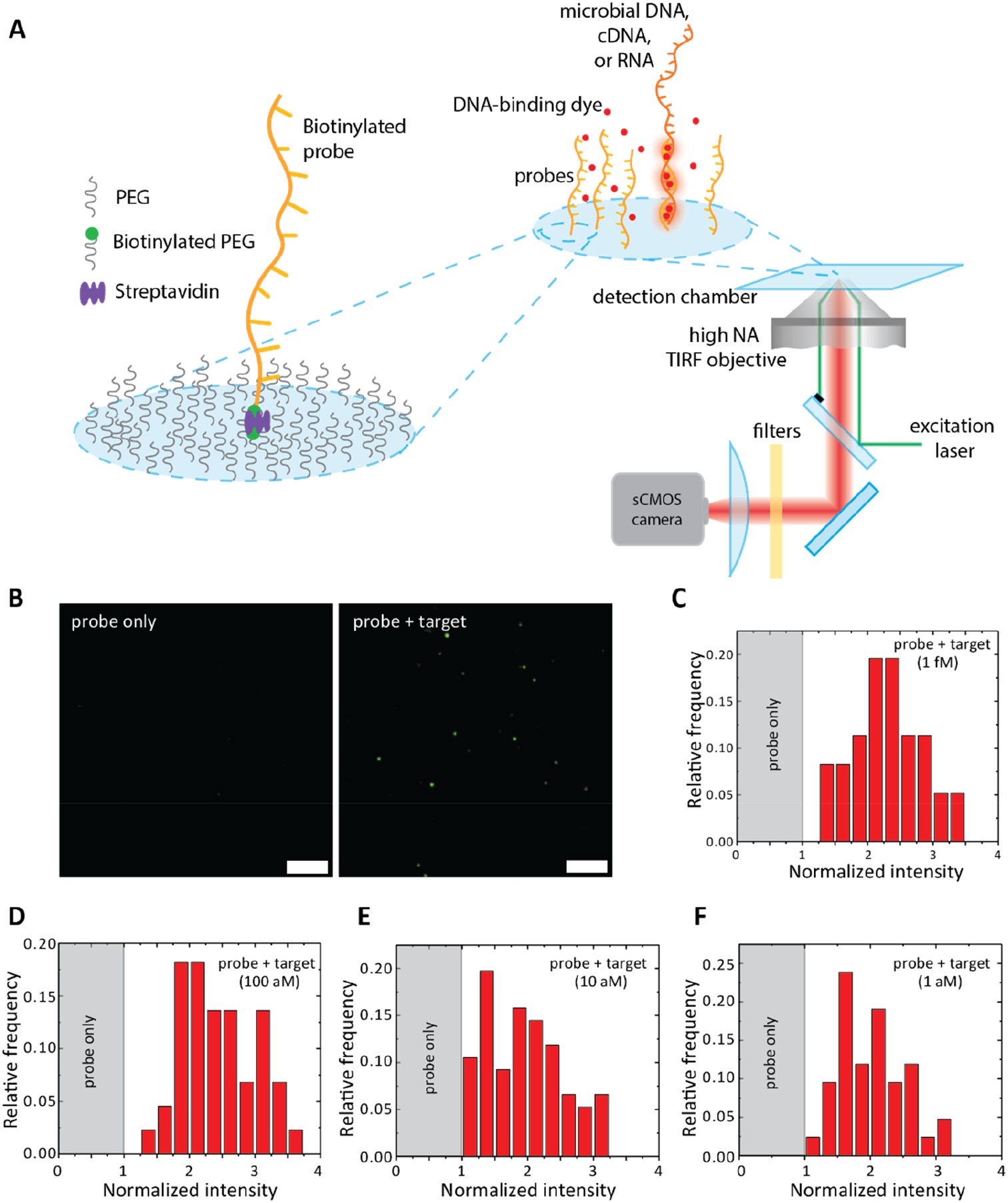
**(A)** Schematic diagram emphasizing the concept of detection.The target microbial sequence hybridizes with the substrate-bound specific probes and recruits a DNA-binding dye making the hybridized sequence brighter. The substrate is passivated with PEG and other surfactants to inhibit non-specific binding of other molecules. A small fraction of the PEG is conjugated to biotin which recruits the biotin-labelled probes through a streptavidin bridge. Light emitted from individual molecules is detected on a camera. **(B)** Images of a substrate to which the single-stranded probe is attached in the presence of the DNA-binding dye only (left) and the 50-mer DNA sequence and the dye (right). Both images have the same intensity scale and the substrate is incubated with a 1fM concentration of the 50-mer DNA target sequence. Scale bars represent 5μm. **(C)** Histograms of intensities of individual spots from samples where the 50-mer DNA are incubated with the probes at concentrations of 1fM **(D)** 100 aM **(E)** 10 aM **(F)** 1 aM. At 1 aM, there are as few as ~600 molecules in a volume of 1ml.

To validate the technique, we hybridized biotinylated 50-base ssDNA probes with complementary 50-base target DNA sequences in a hybridization buffer (see Methods), followed by controlled heating and cooling. Bulk fluorescence measurements using SYBR Safe demonstrated a >3-fold fluorescence increase for 10 nM DNA:DNA hybrids compared to 10 nM ssDNA probes alone (**Figure S2**), confirming the efficacy of intercalators in reporting successful hybridization. The hybridized samples were then incubated in the functionalized sample chamber, while control chambers contained only ssDNA probes. We incubated varying concentrations of target nucleic acid (50 bases) with the corresponding ssDNA probe (50 bases) from picomolar to attomolar levels. For ultra-low concentrations (attomolar), an incubation period of 1–3 hours was required, whereas higher concentrations (picomolar) needed only 30–60 minutes. Multiple images were acquired for each condition, leveraging the tiling feature of our motorized microscope to expedite data collection. After background subtraction, fluorescence intensities of individual spots in the presence of the target nucleic acid were measured and compared to the control (**Figures 1B-F**). Our imaging results exhibited a fluorescence enhancement consistent with bulk measurements, while sample and control were indistinguishable if a mismatched target was used (**Figure S3**), confirming that the method does not produce false-positive signals even at ultra-low concentrations.

Remarkably, Seek-a-Seq enables detection of target nucleic acids at concentrations as low as 1 attomolar (600 molecules per mL) with high specificity (**Figure 1F**). It is worth noting that 1 attomolar is not a fundamental limit of the method, but rather a practical one determined by how much of the incubation chamber is imaged. However, by implanting multiple distinct ssDNA probes in a single chamber designed to target different regions of a microbial genome, and fragmenting the microbial nucleic acids, we anticipate achieving zeptomolar sensitivity, attaining a limit of detection (LOD) unfathomable with amplification-based methods

To demonstrate the ultility and versatility of our approach, we tested whether Seek-a-Seq can detect synthetic SARS-CoV-2 RNA hybridized with complementary 25-base ssDNA probes in the imaging chambers. We observed an >8-fold increase in fluorescence intensity compared to probe-only samples (**Figures 2A-C**), highlighting the robustness of the method. The enhanced signal arises from intercalator binding to the DNA:RNA hybrid regions as well as RNA overhangs and secondary structures within the 5kb synthetic RNA fragment. The SARS-CoV-2 virus was typically present as 10^4^ virions/mL in throat samples taken from individuals several days post initial infection^16^. Any amplification-free detection method should exhibit LODs in the attomolar range to be competitive with NAATs for disease diagnosis. As seen in **Figure 2C**, we could easily achieve LODs better than amplification-based methods without the need for complex sample preparation.

**Figure 2:**
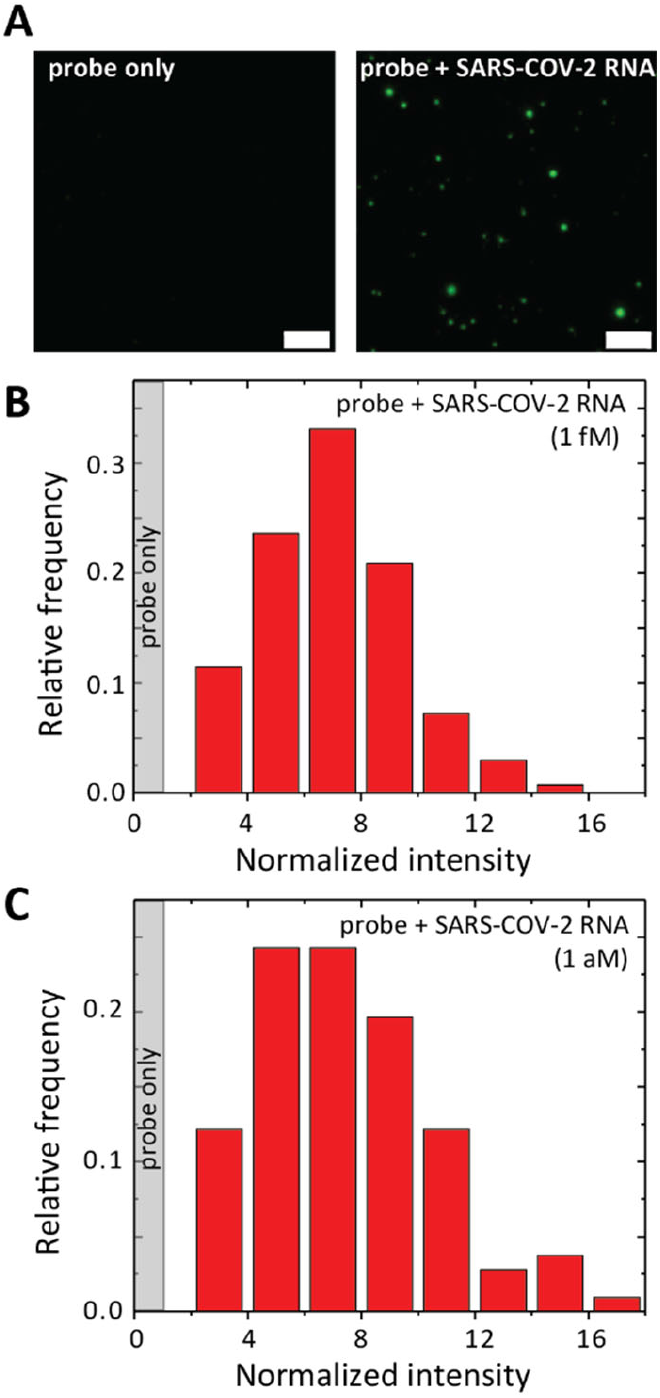
**(A)** Images of a substrate to which the single-stranded probe is attached in the presence of the DNA-binding dye only (left) and the SARS-CoV-2 target sequence and the dye (right). Both images have the same intensity scale and the substrate is incubated with a 1fM concentration of the SARS-CoV-2 RNA. Scale bars represent 5μm. **(B-C)** Histograms of intensities of individual spots from samples where the SARS-CoV-2 RNA was incubated with the 25-mer DNA probe at concentrations of 1fM and 1 aM.

The amplification-free detection strategy and the broad applicability of Seek-a-Seq addresses a significant gap in the nucleic acid diagnostics while overcoming the limitations of existing commercial methods providing an avenue for point-of-care deployment. As the technique is enzyme free, it avoids issues associated with enzyme degradation, storage instability, and batch-to-batch variability, making this approach additionally suitable in settings where low temperature transport and storage is a challenge. Seek-a-Seq integrates single-molecule technologies and advances them into a unique streamlined strategy achieving unprecedented sensitivity exceeding those of traditional NAATs. Taken together, the simplicity, robustness, and cost-effectiveness of our approach position it for widespread adoption in diverse settings, including point-of-care diagnostics, global health crisis management, and resource-limited environments.

## Methods

### Chemicals and reagents

Alconox, Distilled or MilliQ water, ethanol, methanol, potassium hydroxide, acetic acid, 3-aminopropyltriethoxysilane / 3-aminopropyltrimethoxysilane, sodium bicarbonate, mPEG-SVA (MW 5000), biotin-PEG-SVA (MW 5000), streptavidin, biotinylated DNA probes, synthetic SARS-CoV-2 control RNA (Twist Biosciences).

### Surface Passivation and assembly of the detection chamber

#### Cleaning of slides and Coverslips

The microscope slides were scrubbed with Alconox detergent and rinsed thoroughly with distilled water. The coverslips and slides were then placed in Coplin glass staining jars and sonicated with ethanol for 10 mins, followed by rinsing with fresh ethanol solution. Subsequently, the slides and coverslips were rinsed with distilled water to remove any ethanol residue. The water in the jars was replaced with freshly prepared 1M KOH solution for etching. The slides and coverslips were sonicated for 20 to 30 minutes. Thereafter, to remove the KOH traces, the slides were rinsed at least 3 times with distilled water. Clean Nitrogen gas was used to dry the slides and coverslips. During drying, the coverslips were held with Teflon-coated flat-edged tweezers against clean N2 gas. This was followed by plasma cleaning of the coverslips for 15 mins at 30W set at 200 mTorr pressure. Both the slides and coverslips were rinsed with Milli Q water and then dried.

#### Aminosilanization of slides and cover slips

The surface of coverslips was functionalized with the amine group via the aminosilanization chemistry. The amino-silanization solution is prepared by adding 3-aminopropyl triethoxysilane (APTES) to a mixture of methanol and acetic acid (20:1). The cleaned coverslips and slides were rinsed with methanol. Subsequently, the methanol solution in the staining jars was replaced with freshly prepared amino-silanization solution and then incubated for 30 minutes with brief sonication for 1 minute. The aminosilanization reagent was then replaced with methanol and washed 3 times with methanol.

#### Surface PEGylation

The freshly prepared aminosilanized glass surface was passivated by conjugating polyethylene glycol (PEG) succinimidyl valerate at pH 8.5 overnight. Briefly, the amino-silanized slides and coverslips were dried using N2 gas and placed in a humidified chamber with the to-be PEGylated surface facing upwards. The moist environment in the humidified chamber would prevent the drying up of the PEGylation solution. The PEGylation solution was prepared by dissolving methoxy PEG succinimidyl valerate (mPEG-SVA) (250 mg/ml) and biotin-PEG succinimidyl valerate (biotin-PEG SVA) in the ratio 10:1 in freshly-prepared 0.1 M sodium bicarbonate buffer (pH 8.5). The PEGylation solution (~ 100 μl) was placed in between the slide and coverslip and left overnight. The slides and coverslips were carefully disassembled and the PEGylated surface was appropriately labelled, rinsed with distilled water and dried using N2 gas.

#### Assembly of the detection chamber

Circular plexiglass cylinders (1 cm internal diameter, 2 cm height), were cleaned thoroughly and surface-coated with Pluronic F68 polyol. The cylinders were glued to the PEG passivated glass coverslip surface using Epoxy glue. Each detection unit is comprised of two chambers – sample and control. Before sample incubation, the detection chamber was incubated with a streptavidin solution for 30 minutes and washed thrice with buffer.

### Designing probes for SARS-CoV-2

Probes for detecting SARS-CoV-2 were designed for favorable binding to target mRNA sites using the OligoWalk method (ref: Z.J. Lu and D.H. Mathews. (2008) “Efficient siRNA selection using hybridization thermodynamics” Nucleic Acids Research, 36:640-647.). OligoWalk searches for optimal target binding sites in transcripts by considering local RNA structure, probe-target interactions, and probe folding energies. Such considerations help to avoid potential secondary structures that may interfere with probe hybridization and specificity to SARS-CoV-2 compared to other coronaviruses. For 25nt DNA probes, the optimal target binding affinities are about −25 kcal/mol, which are typically found in partially base-paired or unpaired regions.

### Hybridization experiment

The hybridization buffer for DNA: DNA hybridization comprised 10 mM Tris buffer and 100 mM NaCl. In the case of DNA: RNA hybridization, the hybridization buffer consisted of 50% Formamide, 40 mM Piperazine-1, 4-bis (2-ethanesulphonic acid) [PIPES], 400 mM NaCl and 1 mM ethylenediaminetatracetic acid [EDTA] in RNAase/DNAase-free distilled water. A stock solution of 100 μM of the biotinylated DNA probe was diluted to several different concentrations (from nM to aM) and incubated with its complementary DNA strand diluted to the same concentration. The test sample and control were heated at 95°C for 5 mins followed by slow cooling and then incubated in the detection chamber. The probe-only sample was used to normalize the data and report successful hybridization if the intensities in the sample chamber observed were significantly greater than the control chamber.

### Image acquisition and analysis

Samples were imaged either in simple epifluorescence mode or in TIRF mode using an inverted motorized Nikon TIRF system consisting of a Ti stand with perfect focus, 100 × 1.49 NA TIRF objective lens, 488 laser line from the Omicron BrixXHUB ultra-high integrated multimode light engine and Hamamatsu ORCA-Flash4.0 V3 Digital CMOS camera. Samples were imaged in a hybridization buffer containing intercalating dye (SYBRSafe). The microscope control and image acquisition were performed by the NIS Elements software (Nikon Instruments). To ensure homogenous illumination, only the central quarter of the chip of field size 512 × 512 pixels (35 μm X 35 μm) was used for imaging. Tiling (10 × 10) of fields was performed to decrease the overall image acquisition time. The intensity of the fluorescent spots after background subtraction was measured using Fiji (https://imagej.net/software/fiji/).

## Acknowledgements

This work was supported in part by the COVID-19 Facilitator Research Fund (CFR-19) to GTS and KCG. We thank Chandrashekhar U. Murade, Shrutidhara Biswas and Chuan Chang for help with experiments. The research was partially carried out using Core Technology Platform resources at New York University Abu Dhabi.

## Data Availability

All data used in this paper are included in the figures and may be obtained from the corresponding author upon reasonable request.

## Author contributions

GTS conceived the study and directed the research. GTS and AKD designed the experiments. AKD performed most of the experiments, while AS contributed by performing some. AKD analysed the data. GTS and AKD interpreted the findings. HHG and KCG designed some of the probes. AKD wrote the manuscript draft, and GTS and AKD revised subsequent versions. Funding acquisition: GTS and KCG. All authors commented on the manuscript at various stages and have read and agreed to the published version.

## Supporting Information

**Figure S1:**
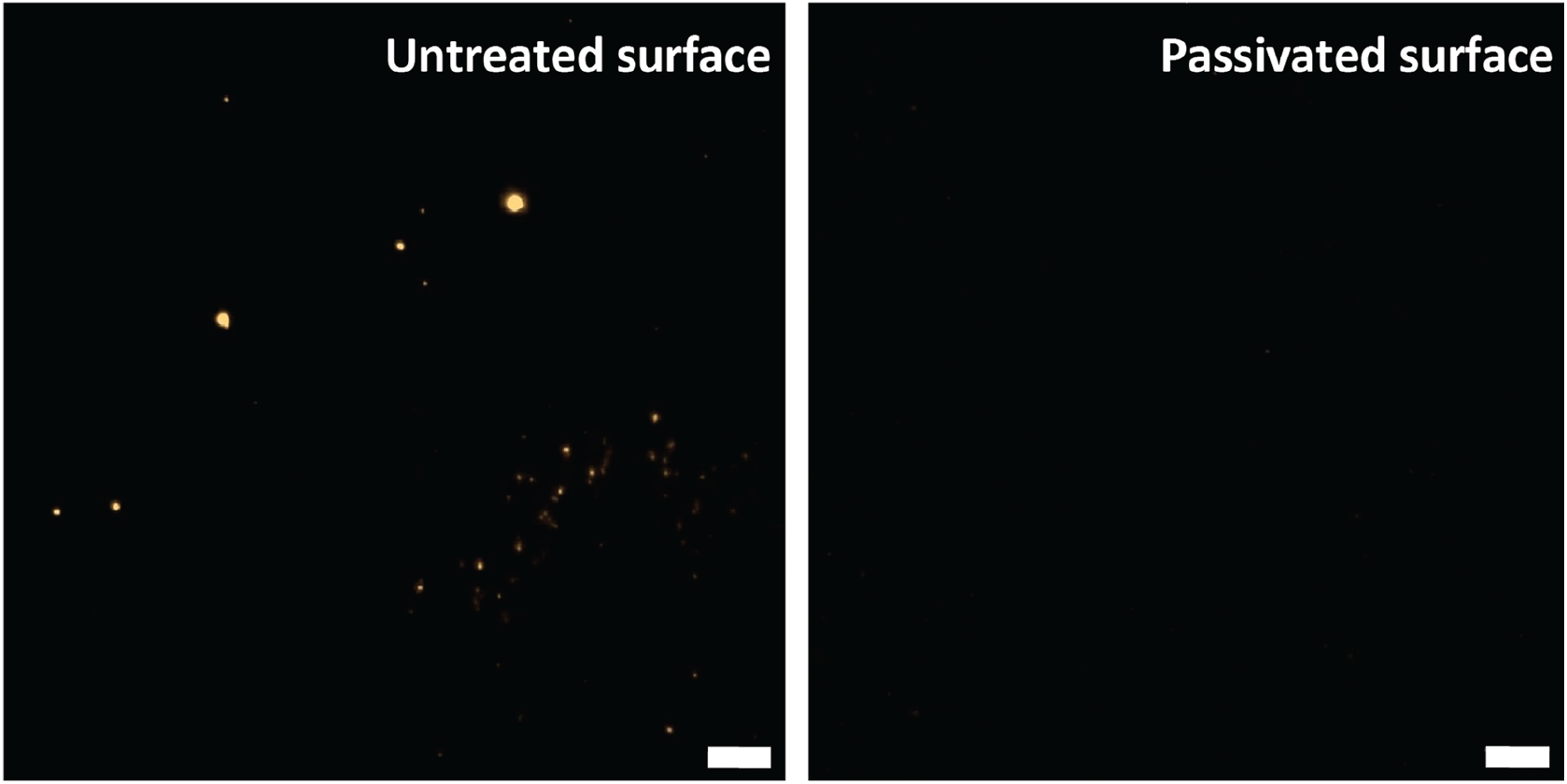
λ–phage DNA was deposited on the surface of untreated (left) and treated (right) detection chambers and visualized in fluorescence using the DNA-binding dye after thorough washing in each case. Both images have the same intensity scale. Scale bars represent 5μm.

**Figure S2:**
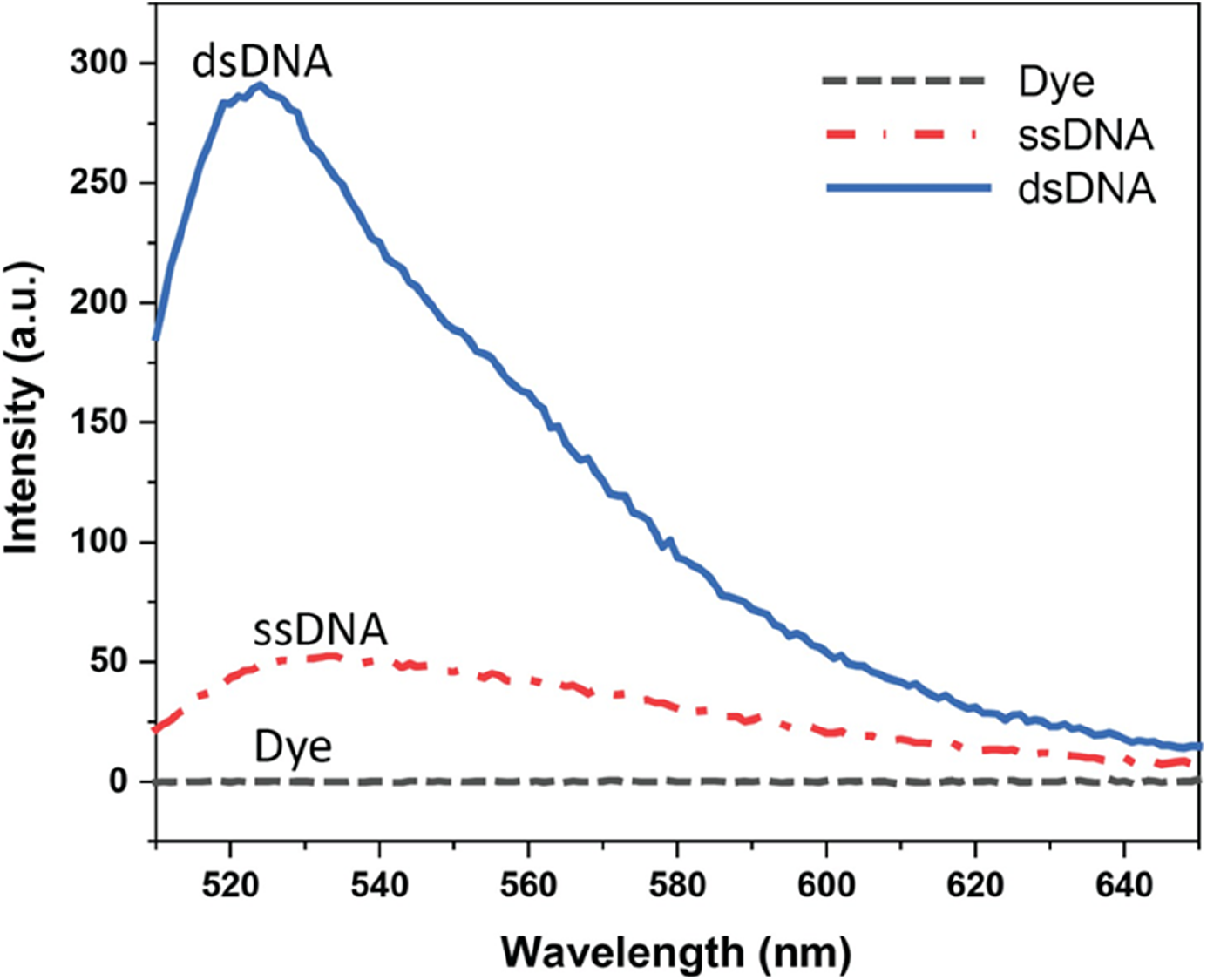
The DNA-binding dye shows enhanced fluorescence when bound to 10 nM double-stranded DNA (dsDNA) when compared to 10 nM single-stranded DNA (ssDNA).

**Figure S3.**
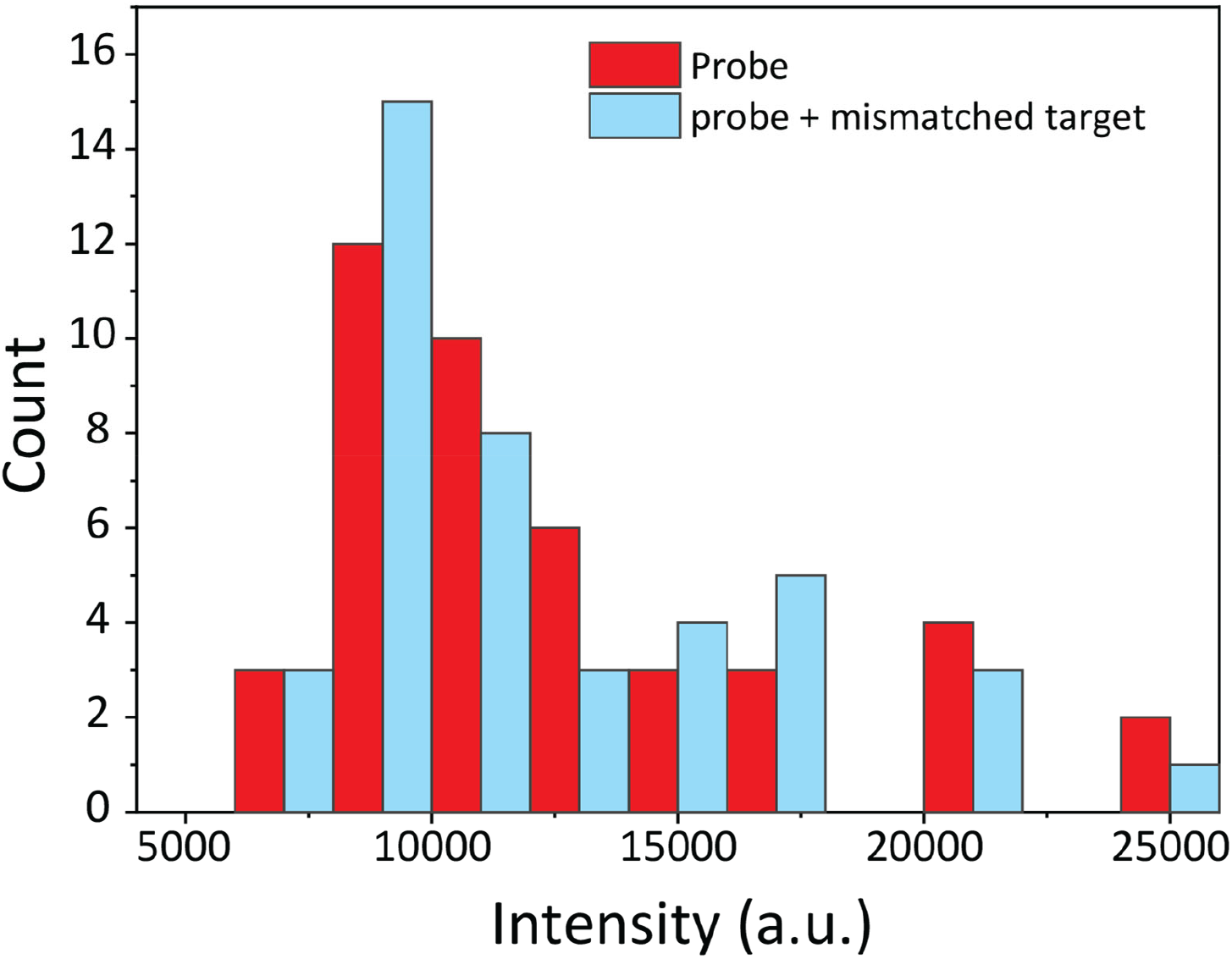
Histograms of intensities of individual fluorescent spots from samples where the probe is attached to the substrate in the presence of the DNA-binding dye only (red) or the dye and a mismatched target (blue). The distributions are indistinguishable emphasizing the specificity of the detection.

## References

1. Kim, T.Y., et al., A review of nucleic acid-based detection methods for foodborne viruses: Sample pretreatment and detection techniques. Food Res Int, 2023. 174(Pt 1): p. 113502.

2. Zhu, Y., et al., Nucleic acid testing of SARS-CoV-2: A review of current methods, challenges, and prospects. Front Microbiol, 2022. 13: p. 1074289.

3. Narasimhan, V., et al., Nucleic Acid Amplification-Based Technologies (NAAT)—Toward Accessible, Autonomous, and Mobile Diagnostics. Advanced Materials Technologies, 2023. 8(20): p. 2300230.

4. Kaminski, M.M., et al., CRISPR-based diagnostics. Nat Biomed Eng, 2021. 5(7): p. 643–656.

5. Cassedy, A., A. Parle-McDermott, and R. O’Kennedy, Virus Detection: A Review of the Current and Emerging Molecular and Immunological Methods. Front Mol Biosci, 2021. 8: p. 637559 (2021)

6. Gootenberg, J.S., et al., Nucleic acid detection with CRISPR-Cas13a/C2c2. Science,. 356(6336): p. 438–442 (2017).

7. Azhar, M., et al., Rapid and accurate nucleobase detection using FnCas9 and its application in COVID-19 diagnosis. Biosens Bioelectron, 183: p. 113207 (2021)

8. de Puig, H., et al., Minimally instrumented SHERLOCK (miSHERLOCK) for CRISPR-based point-of-care diagnosis of SARS-CoV-2 and emerging variants. Sci Adv, 7(32) (2021)

9. Broughton, J.P., et al., CRISPR-Cas12-based detection of SARS-CoV-2. Nat Biotechnol, 2020. 38(7): p. 870–874 (2020)

10. Ning, B., et al., A smartphone-read ultrasensitive and quantitative saliva test for COVID-19. Sci Adv, 7(2) (2021)

11. Sorgenfrei, S., et al., Label-free single-molecule detection of DNA-hybridization kinetics with a carbon nanotube field-effect transistor. Nat Nanotechnol, (2): p. 126–32 (2011).

12. Wang, C., et al., Point-of-care diagnostics for infectious diseases: From methods to devices. Nano Today, 2021. 37: p. 101092.

13. Chandradoss, S.D., et al., Surface passivation for single-molecule protein studies. J Vis Exp, 2014(86).

14. Lamichhane, R., et al., Single-molecule FRET of protein-nucleic acid and protein-protein complexes: surface passivation and immobilization. Methods, 2010. 52(2): p. 192–200.

15. Pan, H., et al., A simple procedure to improve the surface passivation for single molecule fluorescence studies. Phys Biol, 2015. 12(4): p. 045006.

16. Y. Pan. et.al., Viral load of SARS-CoV-2 in clinical samples. Lancet Infect. Dis, 20, pp-411–412

